# Kinetochore-fiber lengths are maintained locally but coordinated globally by poles in the mammalian spindle

**DOI:** 10.1101/2022.11.26.517738

**Authors:** Manuela Richter, Lila Neahring, Jinghui Tao, Renaldo Sutanto, Nathan H. Cho, Sophie Dumont

## Abstract

At each cell division, nanometer-scale components self-organize to build a micron-scale spindle. In mammalian spindles, microtubule bundles called kinetochore-fibers attach to chromosomes and focus into spindle poles. Despite evidence suggesting that poles can set spindle length, their role remains poorly understood. In fact, many species do not have spindle poles. Here, we probe the pole’s contribution to mammalian spindle length, dynamics, and function by inhibiting dynein to generate spindles whose kinetochore-fibers do not focus into poles, yet maintain a metaphase steady-state length. We find that unfocused kinetochore-fibers have a mean length indistinguishable from control, but a broader length distribution, and reduced length coordination between sisters and neighbors. Further, we show that unfocused kinetochore-fibers, like control, can grow back to their steady-state length if acutely shortened by drug treatment or laser ablation: they recover their length by tuning their end dynamics, albeit slower due to their reduced baseline dynamics. Thus, kinetochore-fiber dynamics are regulated by their length, not just pole-focusing forces. Finally, we show that spindles with unfocused kinetochore-fibers can segregate chromosomes but fail to correctly do so. We propose that mammalian spindle length emerges locally from individual k-fibers while spindle poles globally coordinate k-fibers across space and time.

## Introduction

Living systems use simple, small-scale components to build larger and more complex structures. One such structure is the micron-scale spindle, built from nanometer-scale tubulin molecules. The length of the spindle dictates the distance over which chromosomes segregate in dividing cells, and spindle length is known to scale with cell size during development (Good et al., 2013; Hazel et al., 2013; Lacroix et al., 2018; Rieckhoff et al., 2020; Wühr et al., 2008). Defects in spindle length are linked to impaired chromosome segregation (Goshima et al., 1999), cytokinesis errors (Dechant and Glotzer, 2003), and asymmetric division defects (Dudka et al., 2019; Dumont et al., 2007), and long spindles have been hypothesized to come at an energetic cost (Dumont and Mitchison, 2009b). While we know many proteins that can modulate the spindle’s length (Goshima and Scholey, 2010), how they work together to set spindle length and ensure robust chromosome segregation remains poorly understood. We do not know which aspects of spindle length and dynamics are regulated by global cues at the level of the whole spindle, and which are more locally regulated at the level of its components.

Mammalian spindles are built from a network of microtubules, including discrete bundles of microtubules connecting chromosomes to poles. These bundles, called kinetochore-fibers (k-fibers), are made of many microtubules, some of which directly extend from kinetochores to poles (Kiewisz et al., 2022; McDonald et al., 1992; O’Toole et al., 2020). Poles are the convergence points of k-fiber microtubules and other microtubule minus-ends, and they can also serve as an anchor point for centrosomes, if present, and astral microtubules. In many systems, dynein and other motors work together to focus microtubules into asters and poles (Compton, 1998; Goshima et al., 2005a; Heald et al., 1996; Merdes et al., 1996; Roostalu et al., 2018; So et al., 2022). In mammals, k-fiber microtubules turn over on the order of minutes (Gorbsky and Borisy, 1989), detaching from kinetochores and getting replaced. They also exhibit poleward flux, where k-fiber tubulin moves towards poles, with k-fiber plus-ends on average polymerizing and minus-ends appearing to depolymerize at poles (Mitchison, 1989). Both biochemical factors (Goshima and Scholey, 2010) and mechanical force (Akiyoshi et al., 2010; Dumont and Mitchison, 2009a; Nicklas and Staehly, 1967) are thought to tune k-fiber dynamics at both microtubule ends and thereby tune k-fiber length. Microtubule dynamics regulators with length-dependent activities (Dudka et al., 2019; Mayr et al., 2007; Stumpff et al., 2008; Varga et al., 2006) could in principle give rise to the k-fiber’s length scale, beyond simply tuning length. However, k-fiber architecture and organization vary across species, adding complexity to our understanding of how k-fibers set their length. Some spindles, such as in land plants, do not have focused poles (Yamada and Goshima, 2017), and in many species, spindles are composed of short, tiled microtubules indirectly connecting chromosomes to poles (Brugués et al., 2012; Yang et al., 2007), unlike mammalian k-fibers. Broadly, it remains poorly understood which of the mammalian spindle’s emergent properties—such as length, dynamics, and function—emerge globally from the whole spindle, or locally from k-fibers themselves.

While we know that perturbations that affect spindle pole-to-pole distance also affect k-fiber length, and vice versa (Waters et al., 1996), it is still unclear which sets the other. For example, global forces such as cell confinement pulls on poles, leading to k-fiber elongation by transiently suppressing apparent minus-end depolymerization (Dumont and Mitchison, 2009a), but pole-less k-fibers do not elongate under these forces (Guild et al., 2017). Similarly, locally pulling on a k-fiber with a microneedle causes it to stop depolymerizing at its pole and thus elongate (Long et al., 2020). Since poles serve as a connection point for spindle body microtubules, centrosomes, and astral microtubules, they can in principle help integrate physical and molecular information from within and outside the spindle. Indeed, one proposed model is that force integration at spindle poles sets mammalian k-fiber length and dynamics (Dumont and Mitchison, 2009b).However, focused poles may not be essential for setting spindle length, as species without focused poles (Yamada and Goshima, 2017) can still build spindles and set their length. Similarly, inhibiting dynein unfocuses poles but spindles still form albeit with altered lengths in *Drosophila* (Goshima et al., 2005b) and *Xenopus* (Gaetz and Kapoor, 2004; Heald et al., 1996; Merdes et al., 1996), and without a clear effect on mammalian spindle length (Guild et al., 2017; Howell et al., 2001). Further, it is possible to alter kinetochores and microtubule dynamics to shorten k-fibers without a corresponding decrease in the spindle’s apparent length (DeLuca et al., 2006). The role of the mammalian spindle pole on k-fiber structure, dynamics, and function remains an open question.

Here, we ask which emergent properties of mammalian k-fibers require a focused spindle pole. We inhibit pole-focusing forces and ask how k-fiber length, dynamics, and function change when the spindle reaches an unfocused steady-state. Using live imaging, we find that k-fibers can set their mean length without poles but need poles to homogenize and coordinate their lengths between k-fibers. To test whether unfocused k-fibers can recover their lengths, as control ones do, we acutely shorten them using laser ablation or a microtubule-destabilizing drug and show that they recover their length. They do so by tuning their end dynamics and recover more slowly due to reduced baseline dynamics. Thus, k-fiber length is not simply regulated by global pole-focusing forces, but by local length-based mechanisms. Lastly, we show that while the mammalian spindle can move chromosomes without focused poles, it does so with severe segregation and cytokinesis defects. Together, this work indicates that mammalian spindle poles and pole focusing-forces are not required for k-fiber length establishment and maintenance, but for coordinating spindle structure, dynamics, and function across space and time. We propose that the spindle length scale emerges locally at the level of an individual k-fiber, and that robust, coordinated spindle architecture and function arise globally through spindle poles.

## Results

### Spindle poles coordinate but do not maintain kinetochore-fiber lengths

To test whether k-fiber length is set locally or globally, we generated metaphase spindles without focused poles, but with a steady-state length at metaphase. To do so, we overexpressed the dynactin subunit p50 (dynamitin) in PtK2 mammalian rat kangaroo cells, a system with few chromosomes and clearly resolved individual k-fibers. p50 dissociates the dynactin complex and inhibits the pole-focusing forces of its binding partner, dynein (Echeverri et al., 1996; Howell et al., 2001; Quintyne et al., 1999).

We first imaged unfocused spindle assembly in cells overexpressing p50 using long-term confocal fluorescence live imaging with a wide field of view to capture these rare events. While k-fibers seemed initially focused in these cells, these k-fibers eventually lost their connection to centrosomes and became unfocused, exhibiting a similar phenotype to spindle assembly in some NuMA-disrupted cells (Figure 1A, Figure 1—video 1, 2, Silk et al., 2009). We observed disconnected centrosomes seemingly move around freely in cells with unfocused spindles (Figure 1—video 2, 3). The resulting metaphase spindles were barrel-shaped with bi-oriented chromosomes, and they underwent anaphase after several hours instead of about 30 minutes in control (Figure 1A, Figure 1—video 1, 2). While these spindles had no clear poles, we sometimes observed transient clustering of neighboring k-fibers, likely due to residual pole-focusing forces from other minus-end motors or incomplete dynein inhibition. Their interkinetochore distance was indistinguishable from control, suggesting that k-fibers are still under some tension from other forces (Elting et al., 2017; Kajtez et al., 2016; Maiato et al., 2004; Milas and Tolić, 2016), despite not being connected to poles (Figure 1—figure supplement 1). We hereafter refer to these spindles and k-fibers without distinct poles and with reduced pole-focusing forces as “unfocused”.

**Figure 1.**
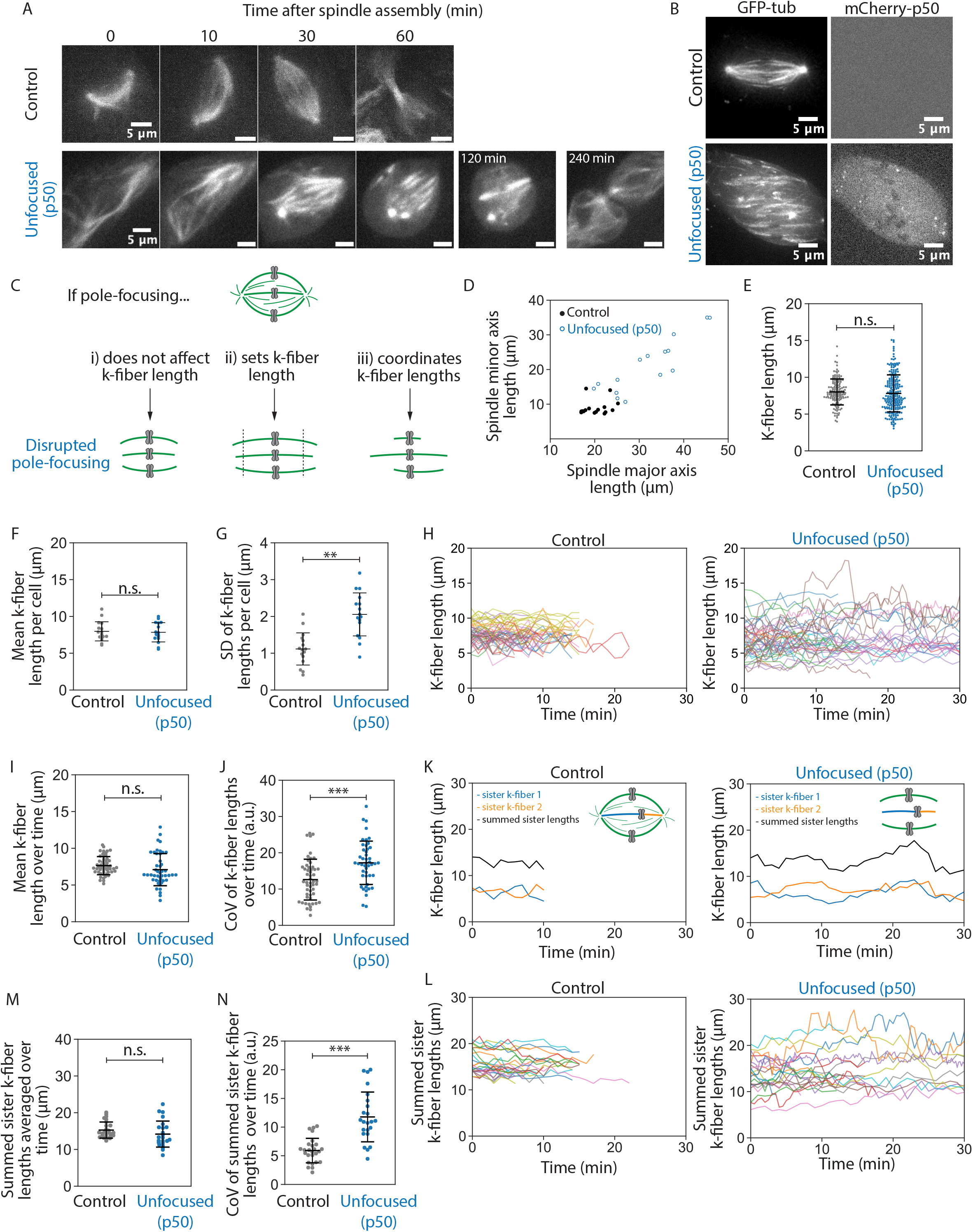
Spindle poles coordinate but do not maintain kinetochore-fiber lengths. See also Figure 1–video 1, 2, 3. **(A)** Representative confocal timelapse images of spindle assembly showing max-intensity z-projections of HaloTag-β-tubulin PtK2 spindles labeled with JF 646, from nuclear envelope breakdown at t= 0 through cytokinesis. mCherry-p50 was infected into unfocused but not control cells. **(B)** Max-intensity z-projections of representative confocal images of PtK2 spindles with GFP-α-tubulin (control and unfocused) and mCherry-p50 (unfocused only). **(C)** Cartoon model of a mammalian spindle with chromosomes (gray) and microtubules (green), with predictions for k-fiber lengths after disrupting poles. Figures D-G are from the same dataset (Control: N = 16 cells; Unfocused: N = 16 cells). **(D)** Spindle major and minor axis lengths in control and unfocused spindles. (Major axis Control = 20.24 ± 2.65 μm, Unfocused = 31.87 ± 7.85 μm, p = 6.3e-5; Minor axis: Control = 8.96 ± 2.12 μm, Unfocused = 21.23 ± 7.61 μm; p = 2.5e-5; Control N = 16, Unfocused N = 15). **(E)** Lengths of control and unfocused k-fibers from z-stacks by live-cell imaging. (Control: n = 144 k-fibers, 8.01 ± 1.76 μm; Unfocused: n = 222 k-fibers, 7.81 ± 2.52 μm; p = 0.38) **(F)** Mean lengths of control and unfocused k-fibers averaged by cell (Control: 7.97 ± 1.30 μm; Unfocused: 7.84 ± 1.31 μm; p = 0.79). **(G)** Length standard deviation of control and unfocused k-fibers per cell. (Control: 1.12 ± 0.44 μm; Unfocused: 2.05 ± 0.58 μm; p = 2.9e-5) Figures H-N are from the same dataset (Control: N = 9 cells, n = 52 k-fibers; Unfocused: N = 9 cells, n = 46 k-fibers) **(H)** Lengths of k-fibers measured over time in control and unfocused spindles. Each trace represents one k-fiber; each color represents a cell. **(I)** K-fiber length averaged over time in control and unfocused spindles. Each point represents one k-fiber. (Control: 7.64 ± 1.23 μm; Unfocused: 7.09 ± 2.19 μm; p = 0.14) **(J)** Coefficients of variation for k-fiber lengths over time in control and unfocused spindles. Each point represents one k-fiber. (Control: 12.60 ± 5.62 a.u.; Unfocused: 17.23 ± 5.98 a.u.; p = 1.8e-4). Figures K-N were analyzed by sister k-fiber pairs (Control: N = 9 cells, n = 26 k-fiber pairs; Unfocused: N = 9 cells, n = 23 k-fiber pairs) **(K)** Lengths of sister k-fibers were measured over time in control and unfocused spindles. One representative k-fiber for each condition is shown in orange, its sister in blue, and their sum in black. **(L)** The sum of sister k-fiber lengths over time in control and unfocused spindles. Each trace is one sister k-fiber pair. **(M)** Summed sister k-fiber lengths averaged over time (from L). Each dot represents one sister k-fiber pair. (Control: 15.27 ± 2.19 a.u. ; Unfocused: 14.18 ± 3.54 a.u.; p = 0.22). **(N)** Coefficient of variation of summed sister k-fiber lengths over time (from L). Each dot represents one sister k-fiber pair. (Control: 5.90 ± 2.14 μm; Unfocused: 11.77 ± 4.34 μm; p = 2.4e-6). Numbers are mean ± standard deviation. Significance values determined by Welch’s two-tailed t-test denoted by n.s. for p≥0.05, * for p<0.05, ** for p<0.005, and *** for p<0.0005.

To measure k-fiber lengths more accurately, we imaged control and unfocused spindles at metaphase using short-term confocal fluorescence live imaging at higher spatial resolution (Figure 1B). If poles do not contribute to k-fiber length, we expect no change in k-fiber length distributions in unfocused spindles (Figure 1Ci). If poles are required to set spindle length, we expect k-fibers with a different mean length in unfocused spindles (Figure 1Cii). If poles merely coordinate lengths, we expect k-fibers with a greater variability of lengths in p50 spindles, but the same mean length (Figure 1Ciii). We first observed that in unfocused spindles, k-fibers were more spread out in the cell, with spindles covering a larger area compared to control and wider spindles tending to be longer (Figure 1D). This is consistent with pole-focusing forces providing contractile forces to compact the spindle (Hueschen et al., 2019). Next, we measured k-fiber lengths in 3D. For control spindles whose k-fibers end at centrosomes at this resolution, we subtracted the radius of the centrosome (0.97 ± 0.10 μm) from the region of measured tubulin intensity (Figure 1—figure supplement 2). Mean k-fiber length in an unfocused spindle (7.81 ± 2.52 μm) was not significantly different than control (8.01 ± 1.76 μm) (Figure 1E). Thus, k-fibers do not require a pole connection to keep their mean length. However, these unfocused spindles showed a greater standard deviation in lengths, so we compared average k-fiber lengths per cell to account for cell-to-cell variability: the mean k-fiber length within each cell was indistinguishable between control and unfocused cells (Figure 1F), but the standard deviation was significantly greater in unfocused cells (Figure 1G). This indicates that spindle poles act to synchronize lengths between neighbors within a spindle, rather than to set and keep length. K-fibers can maintain their average length without poles, but they do so with a greater length variability.

In principle, this greater k-fiber length variability in unfocused spindles could not only come from greater length variability between k-fibers in a given cell (Figure 1G), but also from greater variability over time for each k-fiber. To test this idea, we measured k-fiber lengths over time (Figure 1H, Figure 1—video 3). We observed indistinguishable mean lengths averaged over time in unfocused and control k-fibers and a greater coefficient of variation in unfocused k-fiber lengths over time compared to control (Figure 1I, J). Thus, while unfocused k-fibers still establish and maintain their mean lengths at a similar length scale (Figure 1F, I), their lengths are more variable within a cell (Figure 1G) and over time (Figure 1J) compared to control.

Finally, to test the role of poles in coordinating lengths within the spindle, we compared sister k-fiber lengths over time. During chromosome oscillations, sister k-fiber lengths are normally anti-correlated (Wan et al., 2012). Indeed, in control cells we observed that as one sister k-fiber shortened, the other elongated to maintain a constant sum of their lengths. However, this was not observed in unfocused spindles (Figure 1K). In unfocused spindles, the sum of sister k-fiber lengths was indistinguishable from control when averaged over time, but their sum was less conserved over time, yielding higher coefficients of variation (Figure 1K-N). Thus, poles help coordinate lengths across sister k-fibers such that chromosomes can move within the metaphase spindle while maintaining spindle length.

Together, our findings indicate that spindle poles are not required to globally maintain k-fiber length. Instead, individual k-fibers can locally maintain their length scale over time, and poles and global pole-focusing forces are needed to coordinate k-fiber lengths within the cell and across sister k-fibers, organizing the spindle’s structure in space and time.

### Kinetochore-fibers recover their lengths without focused poles

We have shown that k-fibers can establish and maintain their length independently of poles and pole-focusing forces, but cannot properly organize their lengths within the spindle across space and time. While unfocused k-fibers within a cell maintain their average length over time, we sought to determine whether they can recover their length without focused poles, that is, whether they actively adjust and recover their length if shortened below their steady-state length. First, we used laser ablation to acutely cut and shorten k-fibers and then imaged their regrowth compared to unablated k-fibers (Figure 2A-D, Figure 2—video 1). Due to not capturing the full length of k-fibers in a single z-plane while imaging ablations, we observed a shorter mean length than expected in unfocused unablated k-fibers (Figure 2D); indeed, length analysis of full z-stacks from unablated spindles before ablation yielded an indistinguishable mean k-fiber length from control k-fibers in Figure 1E (Figure 2—figure supplement 1). Ablation generates new microtubule minus-ends on the shortened k-fiber stub, which recruit NuMA and dynein to reincorporate them back into the pole in control cells (Elting et al., 2014; Sikirzhytski et al., 2014). As expected, control ablated k-fibers were transported towards poles and did so while growing back rapidly following ablation, at 0.85 ± 0.09 μm/min on average in the first 5 minutes (Figure 2E). Unfocused k-fibers also grew back, though more slowly at 0.38 ± 0.42 μm/min on average (Figure 2E). They took longer to grow back to the mean length of unablated neighbor k-fibers (Figure 2F). Thus, focused poles and pole-focusing forces are not required for k-fibers to recover their lengths, but are required for rapid length recovery. The latter is consistent with the idea that force on k-fiber ends favors k-fiber growth (Dumont and Mitchison, 2009a; Long et al., 2020; Nicklas and Staehly, 1967). Ultimately, k-fibers can adapt to length changes and maintain a steady-state length locally, without poles.

**Figure 2.**
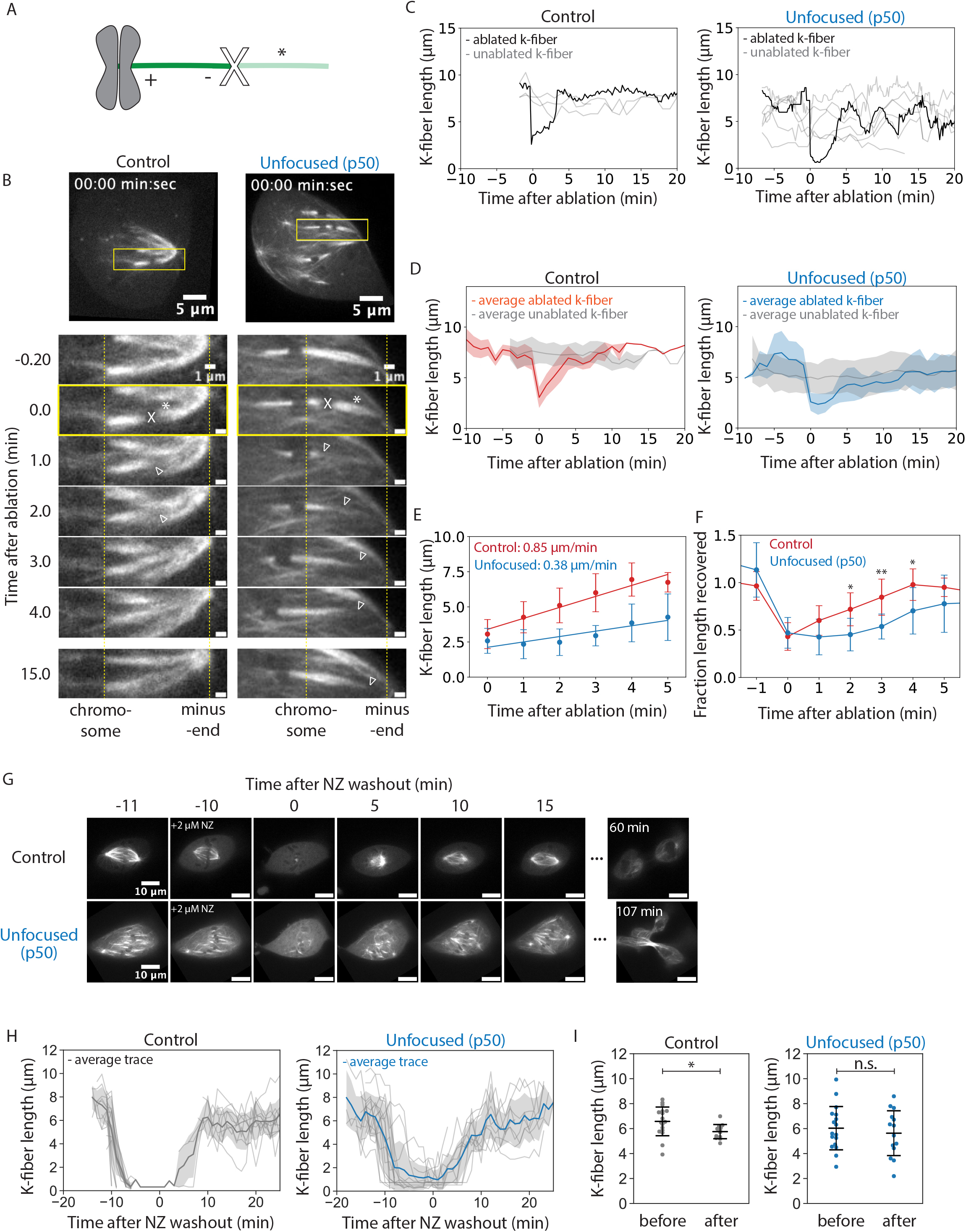
Kinetochore-fibers recover their lengths without focused poles. See also Figure 2–video 1, 2. **(A)** Schematic of a k-fiber after ablation at position X. The k-fiber stub still attached to the chromosome persists with a new minus-end (dark green). The k-fiber segment closer to the pole with a new plus-end depolymerizes away (light green, *). **(B)** Representative confocal timelapse images of PtK2 k-fibers with GFP-α-tubulin and mCherry-p50 (in unfocused only). K-fibers were laser-ablated at t = 0 (X) and followed over time. Empty arrowheads mark newly created minus-ends. Yellow dashed lines are fiduciary marks for the plus- and minus-ends at ablation. **(C)** K-fiber lengths over time in a representative control and unfocused spindle. Gray traces represent unablated k-fibers. The ablated k-fiber is plotted in black. **(D)** Binned and averaged k-fiber lengths over time for ablated control and unfocused spindles. The average length of non-ablated k-fibers is plotted in gray, the average of ablated k-fibers in red for control and blue for unfocused. Shaded colors indicate ±1 standard deviation for their respective condition. (Control: N = 7 cells, n = 8 ablated k-fibers, m = 26 non-ablated k-fibers; Unfocused: N = 6 cells, n = 8 ablated k-fibers, m = 31 non-ablated k-fibers). **(E)** Average growth rates of k-fibers immediately following ablation. Linear regression was performed on binned k-fiber lengths during the first five minutes following ablation (Control: 0.85 ± 0.09 μm/min, Unfocused: 0.38 ± 0.42 μm/min, p = 0.023). **(F)** Fraction of length recovered following ablation relative to the mean of unablated k-fibers in control and unfocused k-fibers. The average trace for unablated k-fibers in D was averaged over time and ablated lengths were normalized to this value. Times with statistically significant differences in length recovery are denoted by *. **(G)** Representative confocal timelapse images of PtK2 spindles with GFP-α-tubulin (in control and unfocused) and mCherry-p50 (in unfocused only), with 2 μM nocodazole added at -10 min and washed out at t = 0. **(H)** Lengths of k-fibers over time during nocodazole washout. All k-fibers are shown with the average trace plotted with ±1 standard deviation shaded in light gray. (Control: N = 3 cells, n = 28 k-fibers; Unfocused: N = 4 cells, n = 23 k-fibers). **(I)** Mean k-fiber lengths before nocodazole and after washout in control and unfocused spindles. (Control before: 6.58 ± 1.15 μm, n = 17; Control after: 5.76 ± 0.57 μm, n = 12, p = 0.02; Unfocused before: 6.03 ± 1.73 μm, n = 17; Unfocused after: 5.63 ± 1.80 μm, n = 14, p = 0.55) Numbers are mean ± standard deviation. Significance values determined by Welch’s two-tailed t-test denoted by * for p<0.05, ** for p<0.005, and *** for p<0.0005.

To test whether neighboring k-fibers or existing microtubule networks provide information for length maintenance, we treated spindles with nocodazole to depolymerize all microtubules, then washed it out and imaged spindle reassembly (Figure 2G, Figure 2—video 2). After 10 minutes, control spindle k-fibers had regrown to within 1 μm of their original length, albeit shorter on average, and unfocused spindle k-fibers fully recovered their average length and grew back into an unfocused state (Figure 2G-I, Figure 2—video 2). Both control and unfocused spindles could enter anaphase after nocodazole washout (Figure 2G, Figure 2—video 2). Thus, cells lacking pole-focusing forces in metaphase can self-assemble unfocused spindles with k-fibers of about the same length as control k-fibers. This supports a model of k-fibers regulating their own lengths without cues from pre-existing microtubule networks or neighboring k-fibers to build a bi-oriented spindle of the correct length scale.

### Kinetochore-fibers exhibit reduced end dynamics in the absence of poles and pole-focusing forces

Given that k-fibers can maintain (Figure 1) and recover (Figure 2) their mean length without poles and pole focusing-forces—albeit regrowing more slowly—we asked whether unfocused k-fibers are dynamic and whether they have reduced dynamics. If dynamics are locally set for each k-fiber, dynamics should not change without poles or pole-focusing forces; if dynamics are set by global pole-focusing forces, we expect different dynamics without poles. In principle, dynamics can be probed using autocorrelation analysis, which reveals the timescale over which k-fibers “remember” their length. If k-fibers were less dynamic and their lengths changed more slowly, this would result in stronger autocorrelation and autocorrelation for a longer period. Indeed, this is what we observed in unfocused k-fibers compared to control, consistent with unfocused k-fibers having reduced dynamics (Figure 3A). We thus sought to measure k-fiber end dynamics and flux.

**Figure 3.**
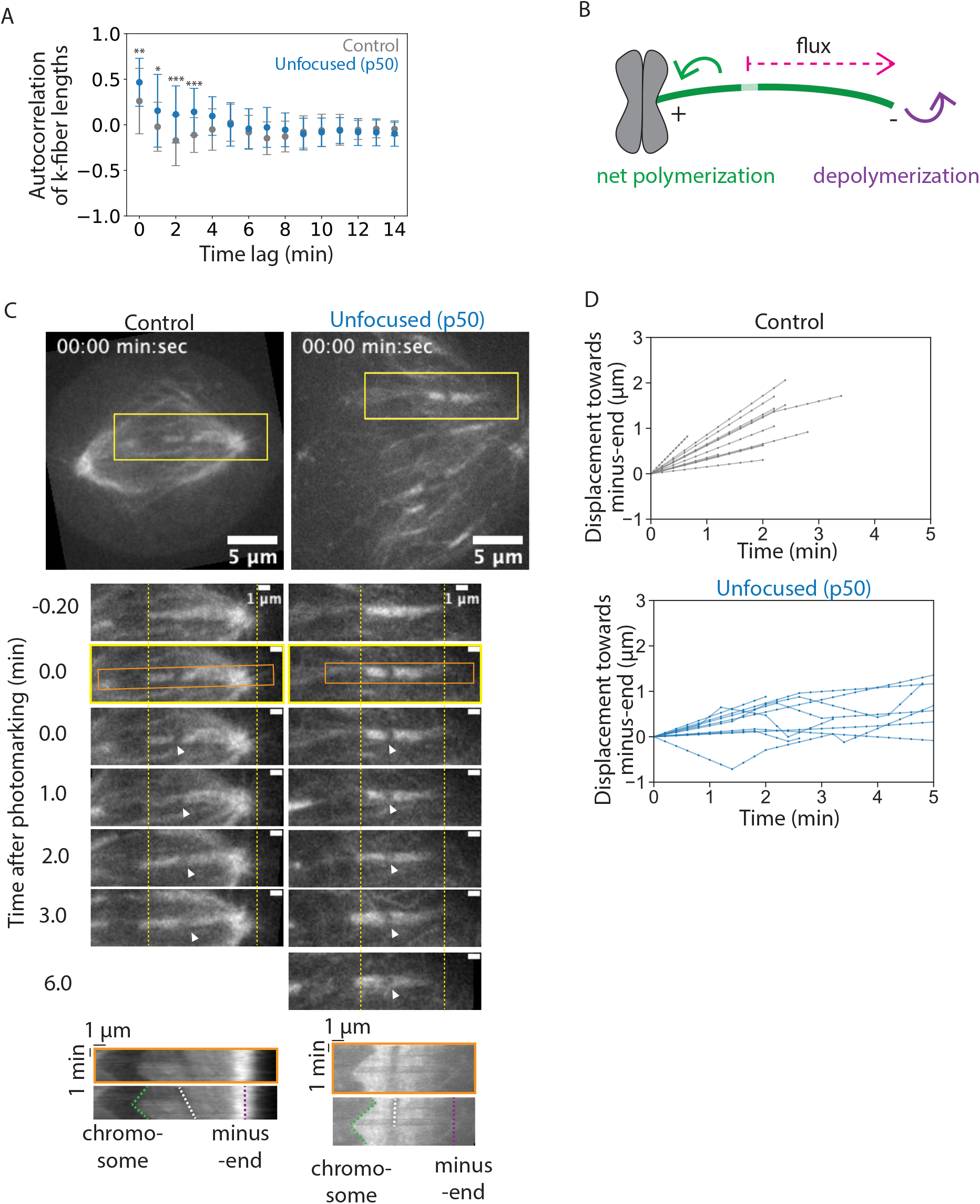
Kinetochore-fibers exhibit reduced end dynamics in the absence of poles and pole-focusing forces. See also Figure 3–video 1. **(A)** Autocorrelation of k-fiber lengths over time from Figure 1H for control and unfocused k-fibers. Calculations and statistical analysis were performed using built-in Mathematica functions, where * indicates p<0.05. **(B)** Schematic of a photomark (light green) on a k-fiber (dark green). The dotted arrow shows the direction the photomark moves with flux in control, where displacement of the mark towards the minus-end increases over time. Net end dynamics are shown by curved arrows (equal at steady-state). **(C)** Representative confocal timelapse images of PtK2 k-fibers with GFP-α-tubulin (in control and unfocused) and mCherry-p50 (in unfocused only). A bleach mark was made at time = 0 and followed over time (arrowhead). Yellow dashed lines are fiduciary marks for the plus- and minus-ends. Below: Kymographs of the above images. Each row of pixels represents a max intensity projection of a 5-pixel high stationary box drawn around the k-fiber at one time point (orange box). **(D)** Minus-end dynamics, measured by displacement of the mark towards the k-fiber’s minus-end over time in control and unfocused k-fibers. Each trace represents one mark on one k-fiber. As the mark fluxes, the distance from the mark to the k-fiber minus-end decreases, and the relative displacement towards the minus-end increases. (Control: N = 8 cells, n = 12 k-fibers; Unfocused: N = 8 cells, n = 11 k-fibers). Numbers are mean ± standard deviation. Significance values determined by Welch’s two-tailed t-test denoted by n.s. for p≥0.05, * for p<0.05, ** for p<0.005, and *** for p<0.0005.

At metaphase, k-fiber ends are dynamic, with poleward flux associating with net polymerization at plus-ends and apparent depolymerization at minus-ends (Mitchison, 1989). Spindle poles have been proposed to regulate minus-end dynamics (Dumont and Mitchison, 2009a; Gaetz and Kapoor, 2004; Ganem and Compton, 2004). To measure k-fiber dynamics, we introduced a bleach mark on a k-fiber and tracked its position over time relative to k-fiber minus-ends (Figure 3B-D, Figure 3—video 1). In control spindles, the mark approached minus-ends at a rate of 0.55 ± 0.29 μm/min, consistent with previous reports (Figure 3D, Figure 4D, Cameron et al., 2006; Mitchison, 1989). In unfocused spindles, the mark approached minus-ends much slower at a rate of 0.13 ± 0.15 μm/min (Figure 3D, Figure 4D). These findings are in contrast to work in *Xenopus* showing that dynein inhibition through p50 overexpression does not impact the flux rate in the central spindle (Yang et al., 2008), but are supported by work in *Xenopus* and in mammals showing that dynein contributes to poleward transport (Burbank et al., 2007; Lecland and Lüders, 2014; Steblyanko et al., 2020). Thus, spindle poles or pole-focusing forces are required for fast k-fiber end dynamics, likely contributing to less efficient k-fiber length maintenance in unfocused spindles.

**Figure 4.**
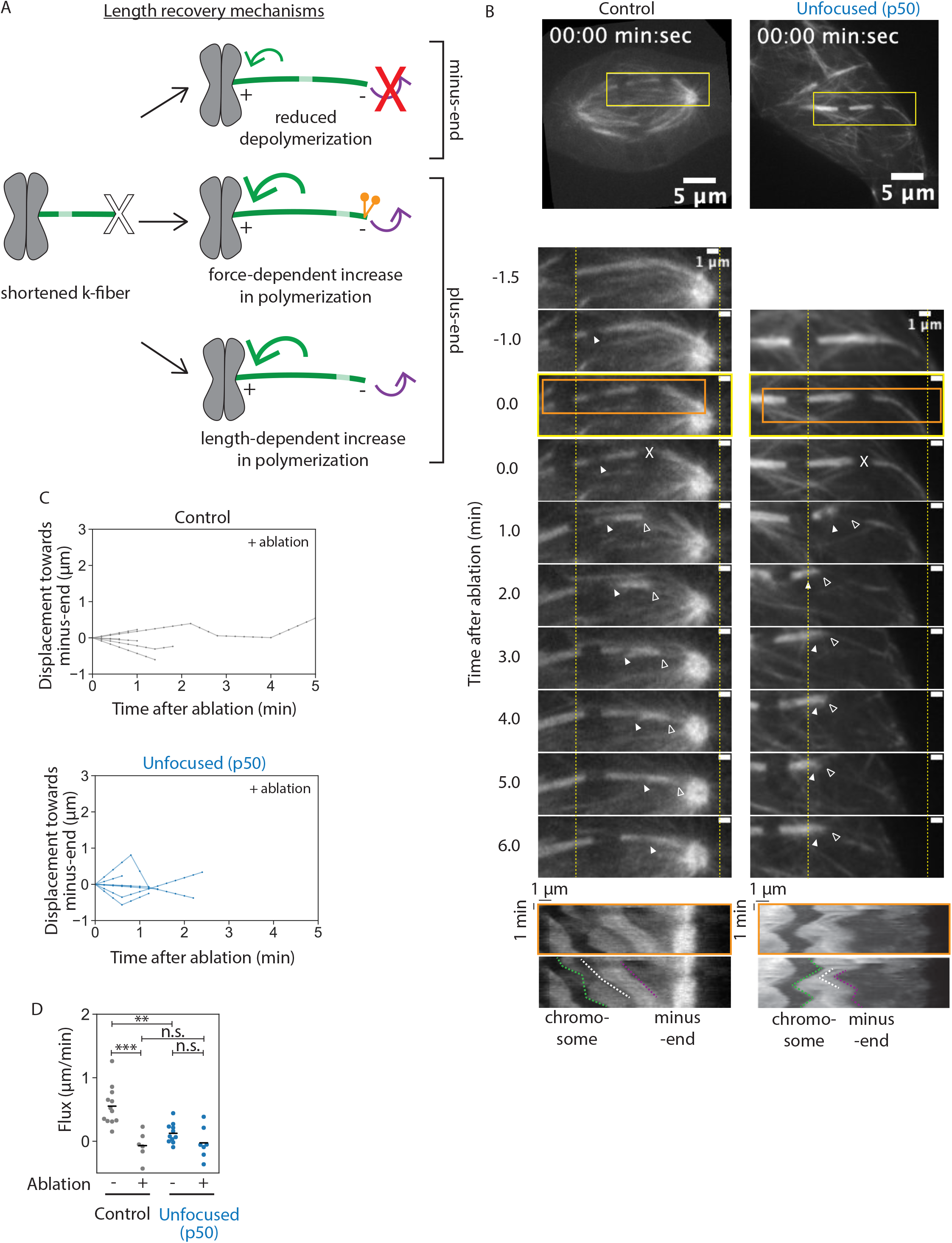
Kinetochore-fibers tune their end dynamics to recover length, without pole-focusing forces. See also Figure 4–video 1. **(A)** Models describing k-fiber length recovery mechanisms. K-fibers shortened by ablation (X) with a photomark (light green) can potentially grow back in different ways: suppression of minus-end depolymerization (top), increased plus-end polymerization induced by forces such as dynein (middle), or increased polymerization in a length-dependent manner (bottom). **(B)** Representative confocal timelapse images of PtK2 k-fibers with GFP-α-tubulin (in control and unfocused) and mCherry-p50 (in unfocused only). Filled arrowhead follows a bleach mark. At t = 0, k-fibers were cut with a pulsed laser at higher power (X). Empty arrowhead follows the new k-fiber minus-end. Yellow dashed lines are fiduciary marks for the plus- and minus-ends. Below: Kymographs of the above images as prepared in Figure 3C. **(C)** Minus-end dynamics were probed by tracking displacement of the mark relative to the k-fiber’s minus-end over time in control and unfocused k-fibers after ablation at t = 0. (Control: N = 5 cells, n = 6 k-fibers; Unfocused: N = 7 cells, n = 7 k-fibers). **(D)** Minus-end dynamics of k-fibers. Rate of photomark displacement towards the minus-end with or without ablation in control and unfocused k-fibers. Each point represents the slope of one trace in Figure 3D or Figure 4C measured by linear regression (Control: mean flux = 0.55 ± 0.29 μm/min, mean flux after ablation = -0.07 ± 0.20 μm/min; Unfocused: mean flux = 0.13 ± 0.15 μm/min, mean flux after ablation = - 0.03 ± 0.23 μm/min; p non-ablated control vs. ablated control = 2.7e-4, p non-ablated control vs. non-ablated unfocused = 5.3e-4, p non-ablated unfocused vs. ablated unfocused = 0.19, p ablated control vs. ablated unfocused = 0.75). Numbers are mean ± standard deviation. Significance values determined by Welch’s two-tailed t-test denoted by n.s. for p≥0.05, * for p<0.05, ** for p<0.005, and *** for p<0.0005.

### Kinetochore-fibers tune their end dynamics to recover length, without pole-focusing forces

The fact that unfocused k-fibers grow back to a steady-state length after being acutely shortened (Figure 2) suggests that they can tune their dynamics after shortening. We thus sought to determine the physical mechanism for length recovery (Figure 4A). One model is that minus-end depolymerization stops or slows—for example, pole-based depolymerization dynamics are lost while k-fiber minus-ends appear separated from the pole (Dumont and Mitchison, 2009a; Long et al., 2020). Another model is that plus-end polymerization increases, which could occur in either a force-dependent manner (Akiyoshi et al., 2010; Dumont and Mitchison, 2009a; Long et al., 2020; Nicklas and Staehly, 1967) or a length-dependent manner (Dudka et al., 2019; Mayr et al., 2007; Stumpff et al., 2008; Varga et al., 2006). Notably, we find that k-fibers can grow back after ablation (Figure 2E) at a rate faster than poleward flux and associated minus-end dynamics in both control and unfocused spindles (0.85 ± 0.09 μm/min vs 0.55 ± 0.29 μm/min in control, 0.38 ± 0.42 vs 0.13 ± 0.15 μm/min in unfocused) (Figure 2E, Figure 4D). Thus, even if minus-end dynamics were suppressed, this would not be sufficient to account for the k-fiber regrowth we observe after ablation, with or without pole-focusing forces.

To directly test how changes in k-fiber length regulate end dynamics, and if this mechanism depends on pole-focusing forces, we ablated a k-fiber and introduced a photobleach mark on it in control and unfocused spindles (Figure 4A, B, Figure 4— video 1). In control spindles, the photomark did not detectably approach the minus-end of the k-fiber during its regrowth (Figure 4B, C), indicating that suppression of minus-end dynamics contributes to k-fiber regrowth, as in *Drosophila* cells (Maiato et al., 2004; Matos et al., 2009). However, while *Drosophila* k-fibers regrow at the rate of poleward flux, these control mammalian k-fibers regrew faster than the rate of flux, indicating that mammalian k-fibers must additionally increase their plus-end dynamics when shortened to reestablish their steady-state length. In unfocused spindles, the photomark also did not detectably approach the minus-end of the k-fiber during its regrowth (Figure 4C), consistent with suppression of any minus-end dynamics, though it was not significantly different from the already slow dynamics and insufficient to account for growth (Figure 4D). Thus, k-fibers can tune their plus-end dynamics to recover their length in the absence of dynein-based pole-focusing forces. This supports a model where k-fiber length is not simply regulated by global pole-focusing forces, but by local length-based mechanisms.

### Spindle poles coordinate chromosome segregation and cytokinesis

So far, we have shown that while a focused pole is not required for setting or maintaining k-fiber lengths (Figure 1, Figure 2), it is required for global spindle coordination (Figure 1) and robust k-fiber dynamics (Figure 3, Figure 4). To test the functional output of focused spindle poles in mammalian cells, we treated control and unfocused spindles with reversine, an MPS1 inhibitor that forces mitotic cells to enter anaphase, even in the absence of dynein activity required for spindle assembly checkpoint satisfaction (Santaguida et al., 2010). Control and unfocused spindles were imaged through anaphase after reversine addition using a single z-plane (Figure 5A, Figure 5—video 1) and also imaged with z-stacks encompassing the whole spindle once before adding reversine, and 20 min after anaphase onset (Figure 5B). In spindles without focused poles, chromatids separated—albeit at twofold reduced velocities compared to control—in the separating chromatid pairs that could be identified (Figure 5C). In the absence of poles or dynein activity, such chromatid separation likely comes from pushing from the spindle center rather than from pulling from the cell cortex (Vukušić et al., 2017; Yu et al., 2019).

**Figure 5.**
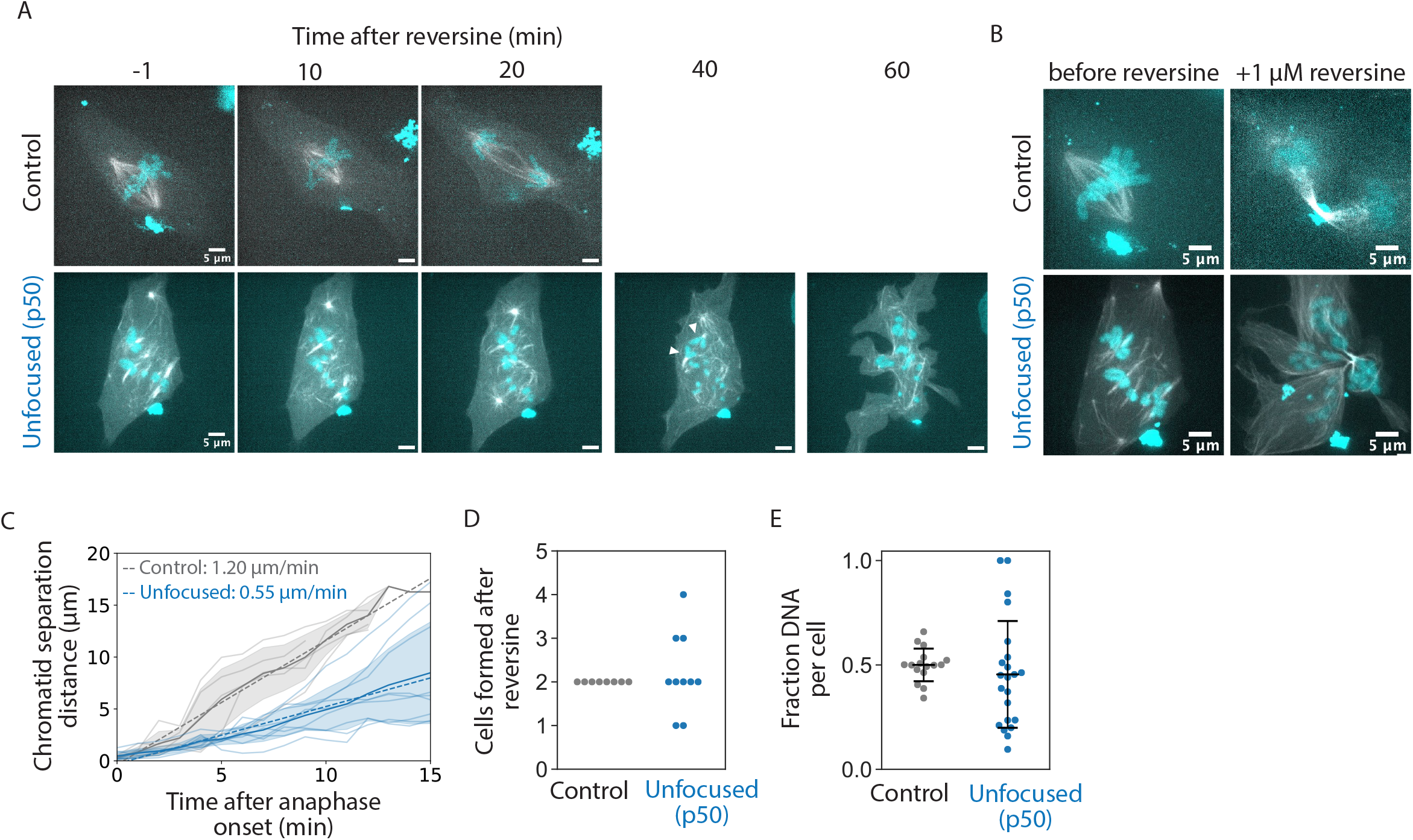
Spindle poles coordinate chromosome segregation and cytokinesis. See also Figure 4–video 1. **(A)** Representative confocal timelapse images of PtK2 spindles with GFP-α-tubulin (in control and unfocused) and mCherry-p50 (in unfocused only) treated with 0.1 or 0.5 μM SiR-DNA with 1 μM reversine added at t = 0. Arrowheads depict an example of sister chromatids separating, later measured in C. **(B)** Max-intensity z-projections before adding reversine and 20 min after anaphase onset for the control and unfocused spindle in A. Figures C-E are from the same dataset. (Control: N = 8 dividing cells; Unfocused: N = 10 dividing cells). **(C)** Sister chromatid separation velocity. For the chromatid pairs that were observed to separate, sister chromatid distance over time was measured for focused and unfocused spindles starting at anaphase onset. Control is plotted in gray, unfocused in blue. Light-colored traces represent one separating chromatid pair, with their average plotted as a dark line with shading representing ±1 standard deviation. The line of best fit for each condition averaged is shown as a dotted line, with their slopes shown. (Control: N = 4 dividing cells, n = 5 chromosome pairs, separation velocity = 1.20 μm/min; Unfocused: N = 3 dividing cells, n = 9 chromatid pairs, separation velocity = 0.55 μm/min). **(D)** Number of “cells” formed after cytokinesis in reversine-treated control and unfocused spindles. (Control: 2 ± 0 cells; Unfocused: 2.20 ± 0.87 “cells”). **(E)** Fraction of chromosome mass per “cell” after reversine treatment. Summed z-projections of chromosome masses were used to calculate the fraction of chromosome mass per cell. (Control: 0.50 ± 0.08 a.u.; Unfocused: 0.45 ± 0.26 a.u.). Numbers are mean ± standard deviation.

However, major segregation and cytokinetic defects were observed in these cells compared to control, consistent with segregation defects observed in k-fibers disconnected from poles (Toorn et al., 2022). Cytokinetic defects and the presence of multiple cytokinetic furrows frequently resulted in the formation of more than two daughter cells in unfocused spindles (Figure 5D). Furthermore, chromosome masses were scattered and unequally distributed in these cells, whereby control daughter cells inherited approximately half of the chromosome mass as measured by DNA intensity, but not daughter cells of unfocused spindles (Figure 5E). Given that focused mammalian spindles lacking dynein pole-focusing forces and lacking Eg5 proceed through anaphase with much milder defects than we observe here (Neahring et al., 2021), we conclude that poles, rather than dynein-based pole-focusing forces, are primarily responsible for these defects. Thus, while many species lack spindle poles, and while unfocused mammalian spindles can still maintain k-fiber length and separate chromatids, spindle poles are essential to coordinate chromosome segregation and cytokinesis in mammalian cells.

## Discussion

Here, we show that in the mammalian spindle, individual k-fibers set and maintain their lengths locally but require the global cue of a focused pole to coordinate their lengths across space and time (Figure 6). Our work reveals that pole-less spindles can set and maintain k-fibers at the same mean length as in control, recovering their steady-state lengths if acutely shortened, but they have impaired dynamics and coordination and are unable to properly segregate chromosomes. We propose a model whereby length is an emergent property of individual k-fibers in the spindle, and where spindle poles ensure that this network of k-fibers is highly dynamic and coordinated across space and time to ultimately cluster chromatids into two future daughter cells.

**Figure 6.**
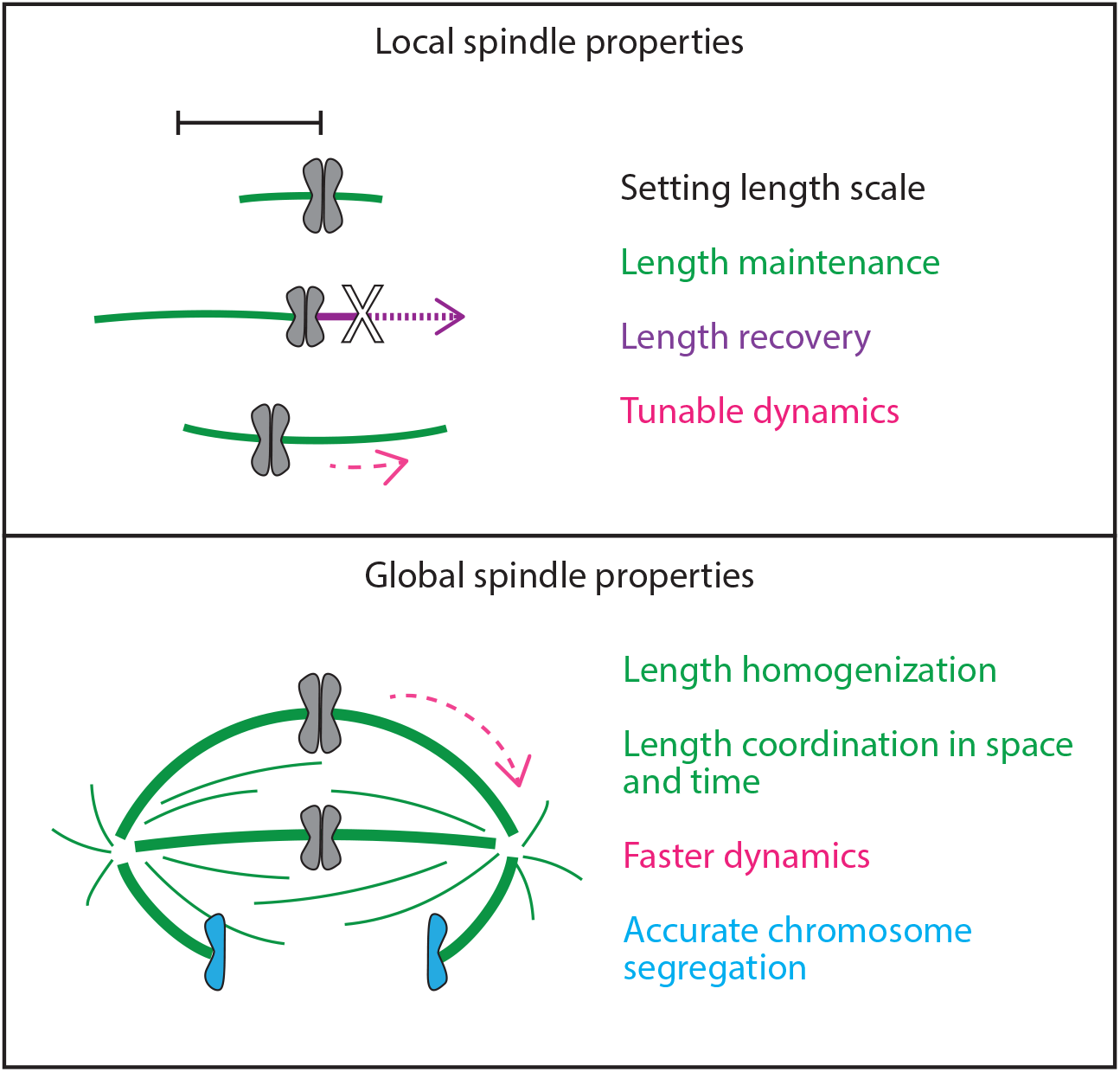
Spindle length is a local spindle property and length coordination is a global spindle property. Cartoon summary of spindle properties set locally versus globally. Setting, maintaining, and recovering length is regulated by individual k-fibers locally, independently of poles and pole-focusing forces. In turn, coordinating lengths across space and time requires global cues from focused poles. In sum, spindle length emerges locally, but spindle coordination emerges globally.

While this work provides insight into k-fiber length establishment and maintenance, what local mechanisms set the k-fiber’s length scale remains an open question. We discuss three models. First, concentration gradients centered on chromosomes (Kalab and Heald, 2008; Wang et al., 2011) could in principle set a distance-dependent activity threshold for spindle proteins that regulate k-fiber dynamics and length. However, it is unclear whether such a gradient with correct length scale and function exists in mammalian spindles. Also, while the globally disorganized structure of unfocused spindles (Figure 1B,D) could lead to modified gradients, the mean length of k-fibers is unchanged (Figure 1E). Second, a lifetime model (Burbank et al., 2007; Conway et al., 2022) stipulates that length is proportional to microtubule lifetime and the velocity of poleward transport, and is sufficient to predict spindle length in spindles with a tiled array of short microtubules. While the length distribution of individual microtubules in unfocused k-fibers is unknown, this model would predict an exponential distribution of microtubule lengths within a k-fiber (Brugués et al., 2012), inconsistent with electron microscopy in control PtK cells (McDonald et al., 1992). Moreover, we observed a more than 4-fold reduced (and near zero) flux velocity in unfocused spindles (Figure 3D), which only a dramatic increase in lifetime could compensate for in this lifetime model. Finally, an “antenna” model (Varga et al., 2006) stipulates that longer k-fibers recruit more microtubule dynamics regulators since they have a longer microtubule antenna to land on. For example, in mammalian spindles, the microtubule depolymerase Kif18A binds k-fibers in a length-dependent way and exhibits length-dependent depolymerase activity, being more active on long k-fibers and thereby shortening them (Mayr et al., 2007; Stumpff et al., 2008). Given that this local antenna model is consistent with our current observations, testing in unfocused spindles whether k-fiber growth rate indeed changes with k-fiber length and testing the role of dynamics regulators in length establishment and maintenance represent important future directions.

Our findings suggest that in response to length changes, k-fibers regulate their plus-end dynamics in an analog manner and their minus-end dynamics in a digital manner. In unfocused spindles, we have shown that the regrowth of shortened k-fibers is driven by an increase in plus-end polymerization, and that this occurs in response to length changes, not simply dynein-based force changes (Figure 4). Consistently, longer k-fibers grow more slowly than shorter ones in a titratable manner in human spindles (Conway et al., 2022). The regulation mechanisms above are all analog in nature. In turn, after ablation, we always observed a near-absence of minus-end dynamics (Figure 4C, D). This is consistent with a switchlike mechanism turning depolymerization on or off, proposed on the basis that tension on k-fibers turns off apparent minus-end depolymerization (Dumont and Mitchison, 2009a; Long et al., 2020). The mechanism behind such digital regulation is not known. One possibility is that a proximal pole structure is required to recruit active microtubule depolymerases, such as Kif2a (Gaetz and Kapoor, 2004; Ganem et al., 2005), to k-fiber minus-ends. In unfocused spindles without a pole, k-fibers would be less dynamic (Figure 3D) based on having fewer depolymerases at their minus-ends. In physical perturbation experiments where k-fibers are separated from the pole center, their apparent minus-end depolymerization would stop (Dumont and Mitchison, 2009a; Long et al., 2020) based on a too-distant depolymerase pool and thus fewer depolymerases at minus-ends. Interestingly, Kif2a can drive spindle scaling in *Xenopus* meiotic spindles (Wilbur and Heald, 2013).

In principle, the concomitant loss of dynein-mediated pole-focusing forces and spindle poles makes it difficult to disentangle the role of each in regulating spindle coordination, maintenance, and function in our findings. However, recent work has revealed that mammalian spindles can achieve similar architecture, exhibit significant— albeit reduced—flux, and segregate chromosomes into two daughter cells whether or not dynein’s recruiter, NuMA, is knocked out (Neahring et al., 2021). This suggests that the severe defects in spindle coordination (Figure 1, Figure 5), dynamics (Figure 3), and maintenance (Figure 2) observed in p50-unfocused spindles are more likely due to the loss of spindle poles than due to the loss of dynein activity per se. Additionally, centrosomes are disconnected from the spindle (Figure 1—video 2, 3), ruling out contributions from centrosomes (Khodjakov et al., 2000) or astral microtubules on k-fiber length regulation at metaphase. Mammalian spindle poles are also required for spindle positioning (Kiyomitsu and Cheeseman, 2012) and have been proposed to help segregate centrosomes (Friedländer and Wahrman, 1970). More work is needed to understand the evolution and function of spindle poles across species and, more broadly, the diversity of spindle architectures across evolution.

We propose that this biological blueprint, where k-fibers locally set and maintain their own length and poles coordinate them globally, robustly builds a complex yet dynamic spindle. For example, we’ve shown that while k-fibers establish their mean lengths locally, global cues homogenize them (Figure 1E, 1G). We put forward the idea that the structural integrity and flexible remodeling of other higher-order structures may also rely on individual parts having all the necessary intrinsic information and self-organization to get the correct linear architecture, with global cues organizing these parts in space and time. More broadly, our work highlights how self-organization at local scales and coordination at global scales can work together to build emergent complex biological structures.

## Figure Legends

**Figure 1—figure supplement 1.**
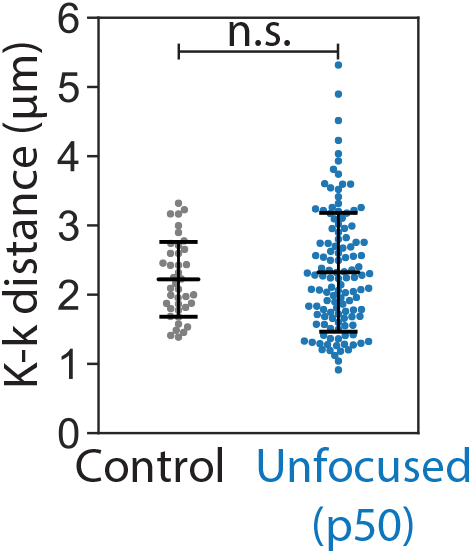
Interkinetochore distance. Interkinetochore distance between sister k-fibers as measured in confocal live-cell imaging of PtK2 spindles expressing GFP-α-tubulin (control and unfocused) and mCherry-p50 (unfocused only). (Control: N = 13 cells, n = 40 kinetochore pairs, 2.22 ± 0.54 μm; Unfocused: N = 16 cells, n = 123 kinetochore pairs, 2.32 ± 0.86 μm; p = 0.38).

**Figure 1—figure supplement 2.**
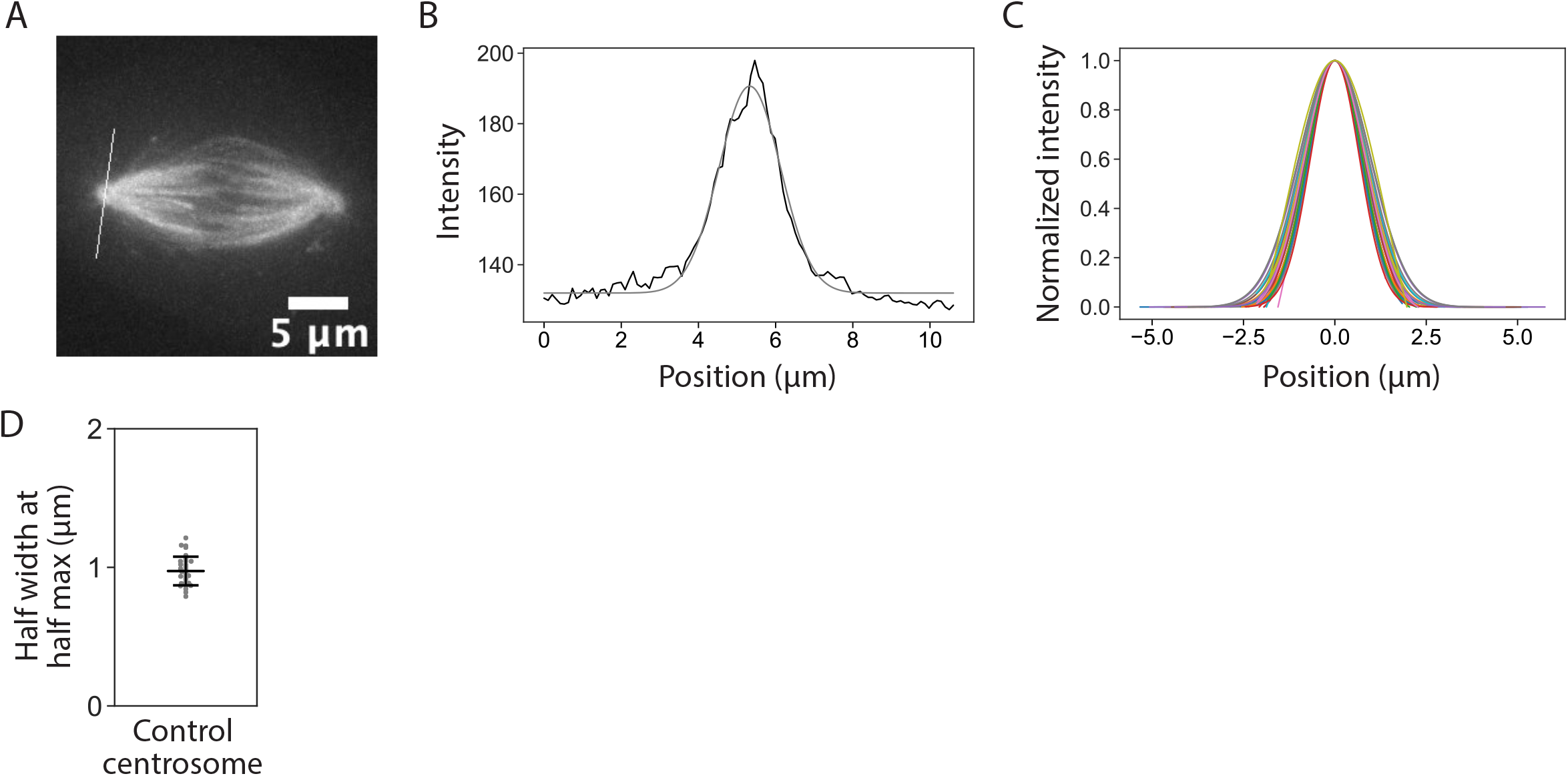
Centrosome radius approximation. **(A)** Example line ROI drawn on a representative centrosome in a max-intensity z-projection of a confocal image of a PtK2 spindle expressing GFP-α-tubulin. **(B)** Line profile of the example centrosome in A. Raw intensity values along the line ROI are plotted in black. These data were smoothed by applying a Gaussian fit and plotted in gray. **(C)** Normalized Gaussian-fitted line profiles of centrosomes. Each color refers to one Gaussian-fitted and normalized centrosome line profile. Traces were normalized by max intensity. **(D)** Centrosome radius was approximated by calculating the half width at half maximum from traces in C. (N = 16 cells, n = 32 centrosomes, 0.97 ± 0.10 μm).

**Figure 2—figure supplement 1.**
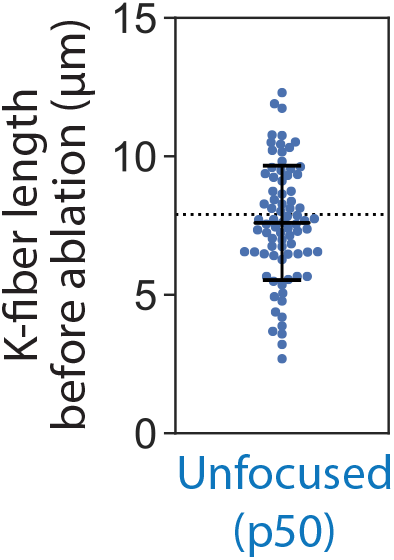
Kinetochore-fiber lengths before ablation. Lengths of k-fibers in unfocused cells prior to ablation. Lengths were measured in 3D from z-stacks of PtK2 cells expressing GFP-α-tubulin and mCherry-p50 taken by confocal live-imaging, as in Figure 1E. The dotted line represents the mean control k-fiber length as calculated in Figure 1E. (N = 4 cells, n = 79 k-fibers, 7.60 ± 2.07 μm).

## Video Legends

Videos are displayed with optimal brightness and contrast for viewing.

**Figure 1—video 1. Control spindle assembly in the presence of pole-focusing forces**. In control cells, k-fibers form focused spindles. See also Figure 1A. Max intensity projection of live confocal imaging of a PtK2 cell expressing HaloTag-tubulin with JF 646 dye. Time is in hr:min with t = 0 at nuclear envelope breakdown. Scale bar, 5μm.

**Figure 1—video 2. Spindle assembly with inhibited pole-focusing forces**. In p50-overexpressing cells, k-fibers grow to eventually form an unfocused spindle. See also Figure 1A. Max intensity projection of live confocal imaging of a PtK2 cell expressing mCherry-p50 and HaloTag-tubulin with JF 646 dye. Time is in hr:min with t = 0 at nuclear envelope breakdown. Scale bar, 5μm.

**Figure 1—video 3. Kinetochore-fiber lengths over time in metaphase: control vs unfocused spindle**. A timelapse of k-fibers in control (left) and unfocused (right) spindles during metaphase. Max intensity projection of live confocal imaging of a PtK2 cell expressing GFP-α-tubulin and mCherry-p50 (unfocused only). Time is in hr:min. Scale bar, 5μm. Videos were cropped and rotated so k-fibers are latitudinal.

**Figure 2—video 1. Ablating kinetochore-fibers: control vs unfocused spindle**. Control (left) and unfocused (right) k-fibers grow back after being severed by a laser. See also Figure 2B. Live confocal imaging of a PtK2 cell expressing GFP-α-tubulin and mCherry-p50 (unfocused only). The ablation site is marked by ‘X’, causing the segment containing the old minus-end of the k-fiber to quickly depolymerize (‘*’). The new stable minus-end is tracked by the empty arrowhead. Time is in min:sec, with ablation occurring at t = 0. Scale bar, 5μm.

**Figure 2—video 2. Spindle assembly after nocodazole washout: control vs unfocused spindle**. Control (left) and unfocused (right) spindles grow back robustly after washing out nocodazole, a microtubule-destabilizing drug. See also Figure 2G. Live confocal imaging of a PtK2 cell expressing GFP-α-tubulin and mCherry-p50 (unfocused only). 2 μM nocodazole was added for 10 min before 10 washes in warmed media were started at t = 0. Time is in hr:min. Scale bar, 5μm.

**Figure 3—video 1. Photobleaching kinetochore-fibers to measure microtubule flux: control vs unfocused spindle**. Control (left) and unfocused (right) k-fibers exhibit poleward flux (reduced in unfocused spindles) as demonstrated by a bleach mark on a k-fiber moving towards a pole over time. See also Figure 3C. Live confocal imaging of a PtK2 cell expressing GFP-α-tubulin and mCherry-p50 (unfocused only). The laser-induced bleach mark is tracked by the filled arrowhead over time as its associated tubulin moves away from the kinetochore towards the minus-end (empty arrowhead). Time is in min:sec, with the photomark created at t = 0. Scale bar, 5μm.

**Figure 4—video 1. Ablating and photomarking kinetochore-fibers: control vs unfocused spindle**. Control (left) and unfocused (right) k-fibers exhibit no measurable minus-end depolymerization during regrowth after ablation. See also Figure 4B. Live confocal imaging of a PtK2 cell expressing GFP-α-tubulin and mCherry-p50 (unfocused only). The ablation site is marked by ‘X’ and the new stable minus-end is tracked by the empty arrowhead. The photomark is tracked by the filled arrowhead and it does not appear to get closer to the other arrowhead at the minus-end over time. Time is in min:sec, with ablation occurring at t = 0. Scale bar, 5μm.

**Figure 5—video 1. A reversine-treated control spindle undergoing anaphase: control vs unfocused spindle**. Control (left) and unfocused (right) spindles treated with a cell cycle checkpoint inhibitor enter anaphase and segregate chromosomes. See also Figure 5A. Live confocal imaging of a PtK2 cell labeled with SiR-DNA (cyan) and expressing GFP-α-tubulin and mCherry-p50 (unfocused only) with 1 μM reversine added. Time is in min:sec, with reversine added at t = 0. Scale bar, 5μm.

## Materials and Methods

### Key Resources Table

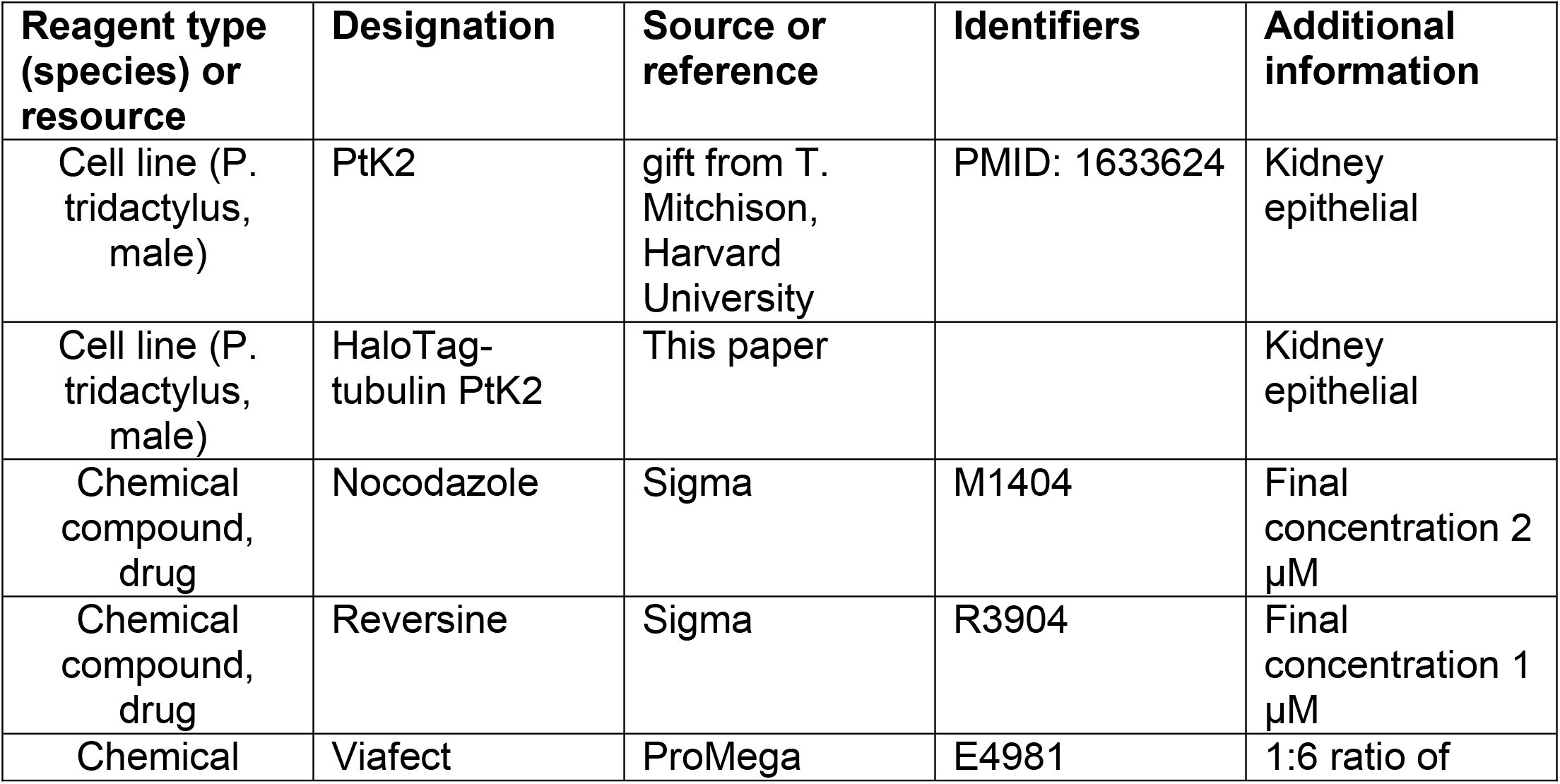
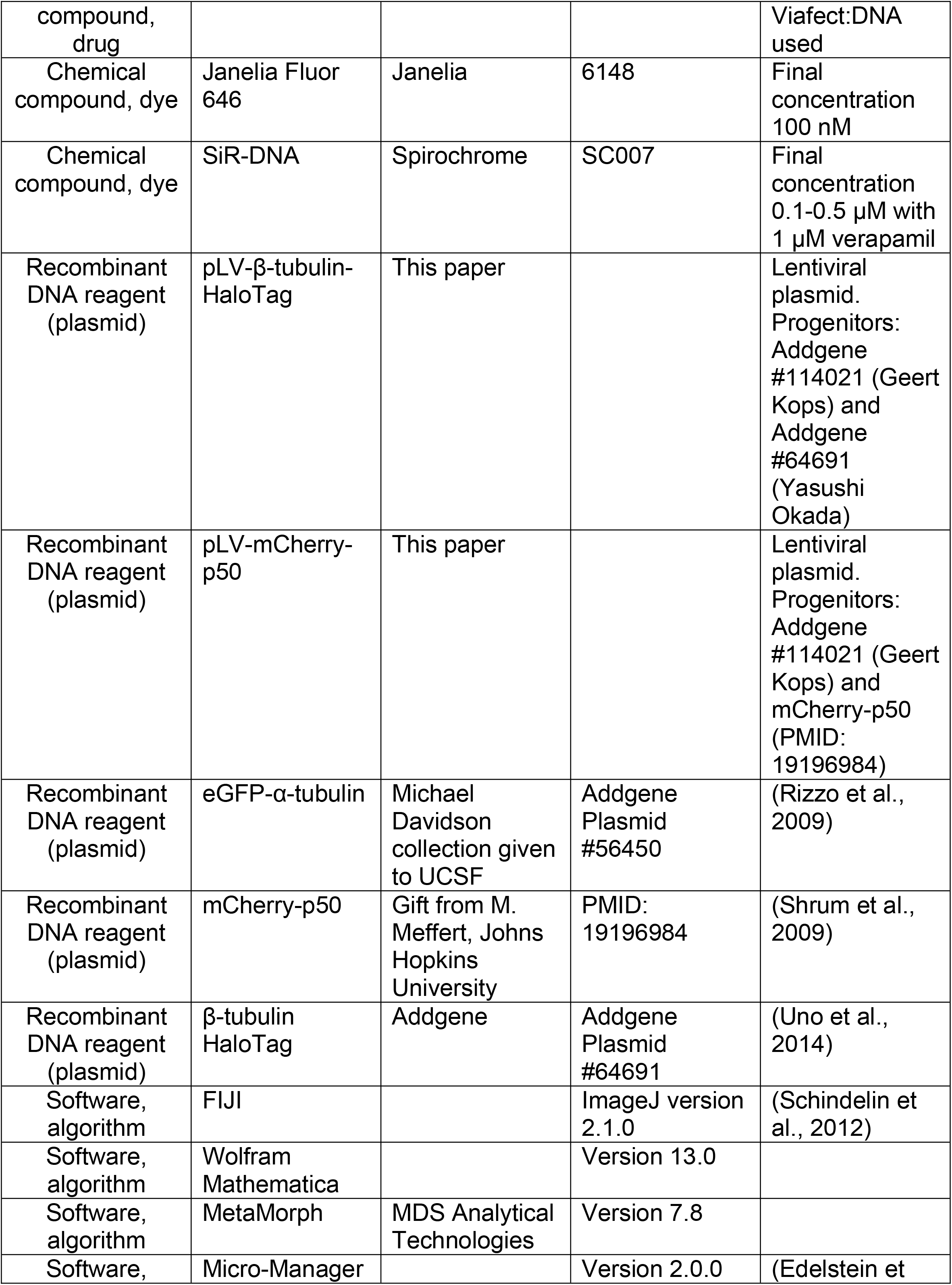
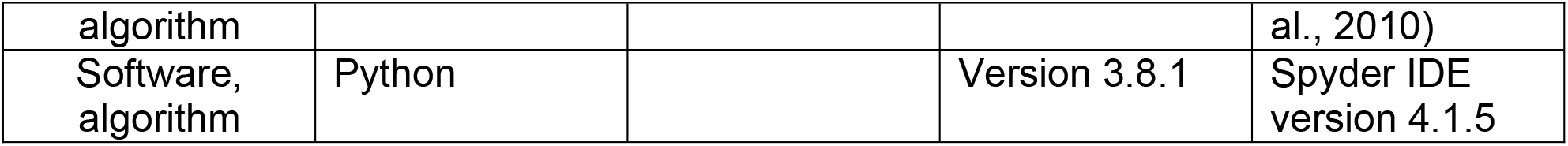

### Cell culture

All work herein was performed using wild-type PtK2 cells (gift from Tim Mitchison, Harvard University). PtK2 cells were cultured in MEM (11095; Thermo Fisher, Waltham, MA) supplemented with sodium pyruvate (11360; Thermo Fisher), non-essential amino acids (11140; Thermo Fisher), penicillin/streptomycin, and 10% heat-inactivated fetal bovine serum (10438; Thermo Fisher). Cells were maintained at 37°C and 5% CO_2_. To visualize microtubules, PtK2 cells were transfected with eGFP-α-tubulin (Clontech) using Viafect (Promega) unless otherwise noted. To inhibit dynein, PtK2 cells were additionally transfected or lentivirally infected with mCherry-p50 (a gift from Mollie Meffert, Johns Hopkins University; Shrum et al., 2009). Transient transfections were prepared in a 100 μl reaction mix per 35 mm dish, including a 1:6 ratio of DNA to Viafect, OptiMEM media up to 100 μl, and eGFP-α-tubulin (0.7 μg) or both eGFP-α-tubulin (0.4 μg) and mCherry-p50 (0.5 μg), and added 3-4 days prior to imaging.

### Lentiviral plasmids and cell line construction

The coding sequences of β-tubulin-HaloTag (Addgene #64691) and mCherry-p50 were cloned into a puromycin-resistant lentiviral vector (Addgene #114021) using Gibson assembly. Lentivirus for each construct was produced in HEK293T cells. To generate the stable polyclonal β-tubulin-HaloTag PtK2 cell line (Figure 1A), wild-type PtK2 cells were infected with β-tubulin-HaloTag virus and selected using 5 μg/ml puromycin. Because p50 overexpression disrupts cell division, mCherry-p50 lentivirus was used to transiently infect each 35mm dish 3-4 days prior to imaging (Figure 1A).

### Imaging

PtK2 cells (gift from T. Mitchison, Harvard University) were plated on 35 mm #1.5 coverslip glass-bottom dishes coated with poly-D-lysine (MatTek, Ashland, MA) and imaged. The cells were maintained at 30-37°C in a stage top incubator (Tokai Hit, Fujinomiya-shi, Japan). Two similar inverted spinning-disk confocal (CSU-X1; Yokogawa Electric Corporation) microscopes (Eclipse TI-E; Nikon) with the following components were used for live-cell imaging: head dichroic Semrock Di01-T405/488/561/647, head dichroic Semrock Di01-T405/488/561, 100x 1.45 Ph3 oil objective, a 60X 1.4 Ph3 oil objective, 488 nm (100, 120, or 150 mW), 561 nm (100 or 150 mW) and 642 (100mW) nm diode lasers, emission filters ET525/36M (Chroma Technology) for GFP, ET630/75M for mCherry, and ET690/50M for JF 646 (Chroma Technology), a perfect focus system (Nikon, Tokyo, Japan), an iXon3 camera (Andor Technology, 105 nm/pixel using 100X objective at bin = 1), and a Zyla 4.2 sCMOS camera (Andor Technology, 65.7 nm/pixel using 100X objective at bin = 1). For imaging, 400 ms exposures were used for phase contrast and 50–100 ms exposures were used for fluorescence. Cells were imaged at 30°C (by default) or 37°C to speed up slower processes (Figure 1A, Figure 2G,H and Figure 5), 5% CO_2_ in a closed, humidity-controlled Tokai Hit PLAM chamber. Cells were imaged via MetaMorph (7.8, MDS Analytical Technologies) or Micro-Manager (2.0.0).

Spindle assembly videos (Figure 1A, Figure 1—videos 1,2) were captured using a 60x objective for a wider field of view, selecting approximately 20 stage positions and imaging overnight at 37°C for 8-10 hours. To capture unfocused spindle assembly, positions containing cells expressing moderate-to-high levels of mCherry-p50 relative to other cells on the dish were selected. Spindles over time were imaged with 1 μm z-slices every minute to avoid photodamage (Figure 1A, Figure 1H). Volumetric spindle images were taken using a 100X objective, with z-slices 0.3 μm apart encompassing the whole spindle (Figure 1B, Figure 5B, Figure 2—supplement 1).

To visualize DNA, 0.1-0.5 μM SiR-DNA (Spirochrome) with 1 μM verapamil were added at least 30 min prior to imaging (Figure 5). To visualize microtubules, 100 nM JF 646 was added to HaloTag-tub PtK2 cells at least 30 min prior to imaging (Figure 1A).

### Photobleaching and laser ablation (Figure 2,3,4)

Photobleaching and laser ablations were performed using 514 or 551 nm ns-pulsed laser light and a galvo-controlled MicroPoint Laser System (Andor, Oxford Instruments) operated through MetaMorph or Micro-Manager. Single z-planes were chosen to pick the clearest k-fiber visible from plus-to minus-end, parallel to the coverslip, that was long enough to ablate. Non-ablated unfocused k-fibers in the same imaging plane were not necessarily parallel to the coverslip, so their full length was not always captured in the single z-plane due to tilt. Photobleaching was performed by firing the laser at the lowest possible power to make a visible bleach mark (∼20% of total power), whereas ablations were performed at the lowest possible power to fully cut a k-fiber (∼60% of total power). K-fiber ablations were verified by observing complete depolymerization of newly created plus-ends, relaxation of interkinetochore distance, or poleward transport of k-fiber stubs (control only). When firing the laser, 1-3 areas around the region of interest were targeted and hit with 5-20 pulses each. Ablations were imaged using one z-plane every 12 s to assay short-term dynamics, then switching to every 1 min after approximately 10 min following ablation to avoid phototoxicity.

### Nocodazole washout (Figure 2)

Z-planes containing the highest number of clearly distinguishable k-fibers, that were parallel to the coverslip, were chosen for imaging. 2 μM nocodazole was swapped into dishes using a transfer pipet while imaging. After 10 min to depolymerize microtubules, dishes were washed 10X in prewarmed media to remove nocodazole and allow spindle reassembly. Spindles were imaged at one z-plane every min to avoid phototoxicity during spindle recovery. To measure k-fiber lengths before nocodazole addition, individual k-fiber traces were averaged over time before drug addition (≤-10 min). K-fiber lengths after drug washout were averaged over time after spindles reached a steady-state length (≥10 min), subtracting centrosome radius for control k-fibers during these times.

### Reversine treatment (Figure 5)

Metaphase spindles were volumetrically imaged with a z-step of 0.3 μm across whole live spindles before reversine addition. The media was then swapped to similar media containing 1 μM reversine and imaged at a single z-plane. 20 min after anaphase onset, cells were again imaged volumetrically as previously described.

### Image analysis

Feature tracking, spindle architecture measurements, and statistical analyses were done in FIJI and Python unless otherwise stated. Videos and images are displayed with optimal brightness and contrast for viewing.

### Spindle major and minor axes length (Figure 1D)

Spindle minor and major axes lengths were determined by cropping, rotating, then thresholding spindle images with the Otsu filter using SciKit.

### K-fiber length (Figure 1,2)

For k-fiber length measurements at a single time point, z-stacks of live spindles were taken with a step size of 0.3 μm across the entire spindle (Figure 1B). Individual k-fibers were measured using a max intensity z-projection of only the slices where that k-fiber was in focus. Line profiles were then measured by drawing ROIs in FIJI with a spline fit line of width 15 pixels, spanning from plus-ends at the start of tubulin intensity next to the chromosome towards minus-ends, using the minimum number of points to recapitulate the curve of the k-fiber. The 3D length was then estimated with the Pythagorean theorem, using the length of the k-fiber’s ROI and the z-height of the slices it spanned (Figure 1E-G). For control k-fibers, the end of the ROI spanning the k-fiber was defined as the center of the pole, and centrosome radius was subtracted to estimate true k-fiber length (Figure 1E-G, I-N, Figure 2C-F, H, I). Since minus-ends of focused k-fibers are not distinguishable in a pole and typically terminate within 2 μm of centrosomes (McDonald et al., 1992), centrosome radius was approximated by drawing line scans through focused poles and measuring the half width at half max intensity. This approximation was used for all subsequent length measurements. For unfocused and ablated k-fibers, minus-ends were defined as the farthest point of visible tubulin intensity corresponding to that k-fiber. Lengths of ROIs were calculated and plotted in Python. K-fiber lengths over time were measured as described above, but from videos with single imaging planes or from max intensity projections based on a step size of 1 μm across the volume of the spindle. K-fiber lengths were then measured using ROIs of width 5 pixels for k-fibers whose plus- and minus-ends were visible across at least 5 frames (k-fiber lengths over time, Figure 1H-N and ablated k-fibers, Figure 2C-F). K-fiber lengths were binned by minute for aggregate analyses. To calculate growth rates for k-fiber lengths over time, linear regression was performed using SciPy on binned k-fiber lengths.

### Tracking photobleach marks along k-fibers (Figure 3, 4)

Spindles of k-fibers with photobleach marks were registered by the tub-GFP channel to account for global spindle translations and rotations. Videos of ablated k-fibers were not registered due to expected translocation of k-fibers stubs after ablation. All videos were trimmed to be isochronous, then flipped, rotated, and cropped so that individual k-fibers with photomarks were latitudinal, with chromosomes on the left and minus-ends on the right. A line with width 5 pixels was drawn along individual k-fibers, and the max intensity projection along the height at each time point was plotted to generate kymographs. Segmented lines were drawn along the kymographs corresponding to the positions of the kinetochore, photomark, and minus-end or pole over time. The distance between the mark and the minus-end over time was calculated and plotted in Python.

### Cell division analysis (Figure 5)

Quantifications of cell division were performed in FIJI. Chromatid separation was quantified by tracking distance between sister chromatids, specifically between the plus-ends of their attached k-fibers, starting the frame before chromatid separation was first observed and ending at the onset of cytokinesis marked by the appearance of a cleavage furrow. To quantify the fraction of chromosome mass per daughter “cell”, “cell” outlines were drawn based on phase contrast images, and the overlap of each cell outline with the summed intensity z-projection of chromosome masses was measured.

### Statistical analysis

Statistical analyses were performed in Python using NumPy and SciPy unless otherwise stated. Linear regression was performed using SciPy. In the text, whenever we state a significant change or difference, the p-value for those comparisons was less than 0.05. In figures, * indicates p<0.05, ** p<0.005, and *** p<0.0005. In the figure legends, we display the exact p-value from every statistical test made. We used a two-tailed Welch’s t-test everywhere unless otherwise stated, since this compares two independent datasets with different standard deviations. Legends include n, the number of individual measurements made, and N, the number of unique cells assayed for each condition.

### Autocorrelation (Figure 3A)

Autocorrelation analysis was performed using Wolfram Mathematica 13.0. The autocorrelation is calculated by the built-in function “CorrelationFunction”. By this definition, the autocorrelation of a k-fiber at lag *h* is 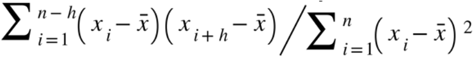 where *x*_*i*_ is k-fiber length at time *i* and 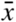 is the mean of *x*_*i*_. The standard deviation is calculated by the built-in function “StandardDeviation”. Statistical significance was performed using the built-in function “LocationTest” at each *h*.

### Script packages

All scripts were written in Python using Spyder through Anaconda unless otherwise stated. Pandas was used for data organization, SciPy for statistical analyses, Matplotlib and seaborn for plotting and data visualization, SciKit for image analysis, and NumPy for general use. FIJI was used for video formatting, intensity quantification, kymograph generation, and tracking k-fibers.

### Video preparation

Videos show a single spinning disk confocal z-slice imaged over time (Figure 2—video 1, Figure 2—video 2, Figure 3—video 1, Figure 4—video 1, Figure 5—video 1) or a maximum intensity projection (Figure 1—video 1, Figure 1—video 2, Figure 1—video 3) and were formatted for publication using FIJI and set to play at 10 fps.

## Acknowledgments

We thank Tim Mitchison for PtK2 cells, Mollie Meffert for the p50 construct, Dan Needleman, Trina Schroer, Wallace Marshall, Orion Weiner, David Agard, Fred Chang, and Christina Hueschen for helpful discussions, and Arthur Molines, Alex Long, Miquel Rosas Salvans, and other members of the Dumont Lab for discussions and critical reading of the manuscript. This work was supported by NIHR35GM136420, NSF CAREER 1554139, NSF 1548297 Center for Cellular Construction, Chan Zuckerberg Biohub, UCSF Byers Award (SD); NSF Graduate Research Fellowship (MR), and the ARCS Foundation (MR); Fannie and John Hertz Foundation Fellowship (LN); American Heart Association Predoctoral Fellowship (NC); and UCSF Discovery Fellows Program (LN, NC).

